# The VapBC Toxin–Antitoxin System Enhances *Shigella flexneri* Fitness Through Coordination of Metabolic Stress Adaptation and Virulence

**DOI:** 10.1101/2025.06.29.662172

**Authors:** Estelle Saifi, Elisabeth Ageron, Keith Egger, Caroline Reisacher, Eric Frapy, Morgan Lamberioux, Shelley Payne, Jost Enninga, Laurence Arbibe

## Abstract

*Shigella flexneri* is a facultative intracellular pathogen that causes bacillary dysentery by invading and replicating within intestinal epithelial cells. Successful intracellular survival requires the bacterium to balance metabolic adaptation with the sustained expression of virulence programs by the type III secretion system (T3SS). Toxin–‘antitoxin (TA) systems, including the highly conserved type II VapBC module, are classically associated with plasmid maintenance via post-segregational killing, but their broader roles during infection remain poorly understood. Here, we investigate the function of the VapBC system during *S. flexneri* infection. We show that the *vapBC* operon is activated in response to intracellular stress and that VapC-dependent cleavage of initiator tRNA*^fMet^* occurs specifically during infection. The stringent response, primarily mediated by SpoT, appears to influence operon responsiveness by maintaining vapB expression levels, while iron limitation emerges as a strong activator of *vapC* transcription and activity. Deletion of *vapBC* impairs T3SS activation and bacterial dissemination, despite normal invasion, and induces a strong host interferon response, including the upregulation of guanylate-binding proteins (GBPs), known to restrict bacterial spread. Transcriptomic profiling of the *ΔvapBC* mutant reveals downregulation of core metabolic genes and upregulation of envelope stress and other TA modules, indicating a loss of intracellular homeostasis. These findings uncover a novel role for VapBC in promoting *Shigella* fitness by coordinating stress adaptation, virulence expression, and immune evasion, thereby sustaining the bacterium’s intracellular lifestyle.

## Introduction

*Shigella flexneri* is a facultative intracellular pathogen responsible for bacillary dysentery, a severe inflammatory disease of the human colon that remains a major public health concern in low- and middle-income countries (Kotloff, 2017). A hallmark of *Shigella* pathogenesis is its ability to invade intestinal epithelial cells, escape the phagocytic vacuole, replicate within the cytosol, and spread to adjacent cells via actin-based motility. These processes are critically dependent on the type III secretion system (T3SS), which injects bacterial effectors into host cells to manipulate host signaling and suppress immune responses (Schnupf & Sansonetti, 2019).

Once inside host cells, *S. flexneri* encounters multiple environmental challenges, including nutrient deprivation, oxidative stress, and cell-autonomous immune defenses (Lucchini et al., 2005)(Pieper et al., 2013)(López-Jiménez et al., 2024). To survive and replicate under these conditions, the bacterium must finely regulate its gene expression to adapt metabolically while maintaining virulence. This coordination is particularly demanding, as both stress adaptation and virulence expression draw on shared and limited cellular resources, forcing the bacterium to carefully balance energy allocation between survival and pathogenesis (Raghunathan et al., 2009). Successfully managing these opposing pressures is essential for intracellular persistence and disease progression.

Toxin–antitoxin (TA) systems, particularly those of type II, are widespread among bacteria, with well-established roles in phage inhibition and in the vertical stabilization of mobile genetic elements through post-segregational killing (PSK), a process in which plasmid-free daughter cells are eliminated and which was only recently confirmed through single-cell analysis (Leroux & Laub, 2022) (Fraikin & Van Melderen, 2024). Over the past decade, a prevailing theory has positioned type II TA systems as key mediators of antibiotic persistence, proposing that they induce growth arrest in subpopulations of “persister” cells, exemplified by *Salmonella enterica* serovar *Typhimurium* within macrophages (Helaine et al., 2014). However, accumulating contradictory evidence has increasingly challenged this model (LeRoux et al., 2020; Maisonneuve et al., 2018). Thus, despite their abundance, the in vivo roles of TA systems, particularly under physiologically relevant infection conditions, remain poorly understood (Jurėnas & Van Melderen, 2020; Pizzolato-Cezar et al., 2023).

Among these systems, the VapBC family is one of the most prevalent type II TA modules in pathogenic bacteria (Pandey & Gerdes, 2005). The VapB antitoxin comprises two domains: an N-terminal DNA-binding domain and a C-terminal intrinsically disordered region that neutralizes the VapC toxin (Loris & Garcia-Pino, 2014). VapC, in turn, harbors a PIN (PilT N-terminal) domain with endoribonuclease activity, capable of cleaving specific cellular RNA targets such as tRNAs and rRNAs (K. Winther et al., 2016; K. S. Winther & Gerdes, 2011). In *Shigella* species, the VapBC module is known to stabilize the large virulence plasmid through PSK (McVicker & Tang, 2016; Sayeed et al., 2000). In vitro studies have shown that VapC toxins in Enterobacteriaceae can specifically cleave initiator tRNA *^fMet^*, leading to translational arrest when overexpressed (K. S. Winther & Gerdes, 2009, 2011). However, the *in vivo* function of VapBC, particularly under conditions encountered during infection, remains largely unexplored.

Here, we investigate the contribution of the VapBC TA system to the intracellular lifestyle of *Shigella flexneri*. Using transcriptomic profiling, tRNA cleavage assays, and infection models, we show that the *vapBC* operon is activated during infection and plays a critical role in modulating bacterial stress responses, virulence gene expression, and evasion of cell-autonomous immunity. These findings reveal a novel function for VapBC in supporting *Shigella*’s intracellular fitness by enabling adaptive reprogramming under host-imposed stress.

## Results

### The operon *vapBC* is activated upon *Shigella flexneri* infection

To uncover the activation of the *vapBC* type II toxin-antitoxin system during *Shigella* infection, we performed dual RNA-sequencing data from infected enterocytic HCT116 cells at 5 hours post-infection). Consistent with its ability to activate host innate immune responses, the analysis showed elevated expression of inflammatory genes in infected-host cells (**Table S1**). Interestingly, in the intracellular pathogen transcriptome, we observed a decrease in read coverage at the tRNA*^fMet^* at the anticodon stem-loop compatible with VapC ribonuclease activity, as described (K. S. Winther & Gerdes, 2011)(**Figure 1a**). Likewise, cleavage at the tRNA*^fMet^* during *Shigella* infection was confirmed by northern blot (**Figure 1b**). This cleavage was not detected when cells were infected with the non-invasive *Shigella* strain BS176, nor in *Shigella* strain maintaining in culture cell medium (referred as “extra-cellular M90T *Shigella* strain”), indicating that it occurs during the intracellular lifestyle of the pathogen (**Figures 1 b-c**).

**Figure 1.**
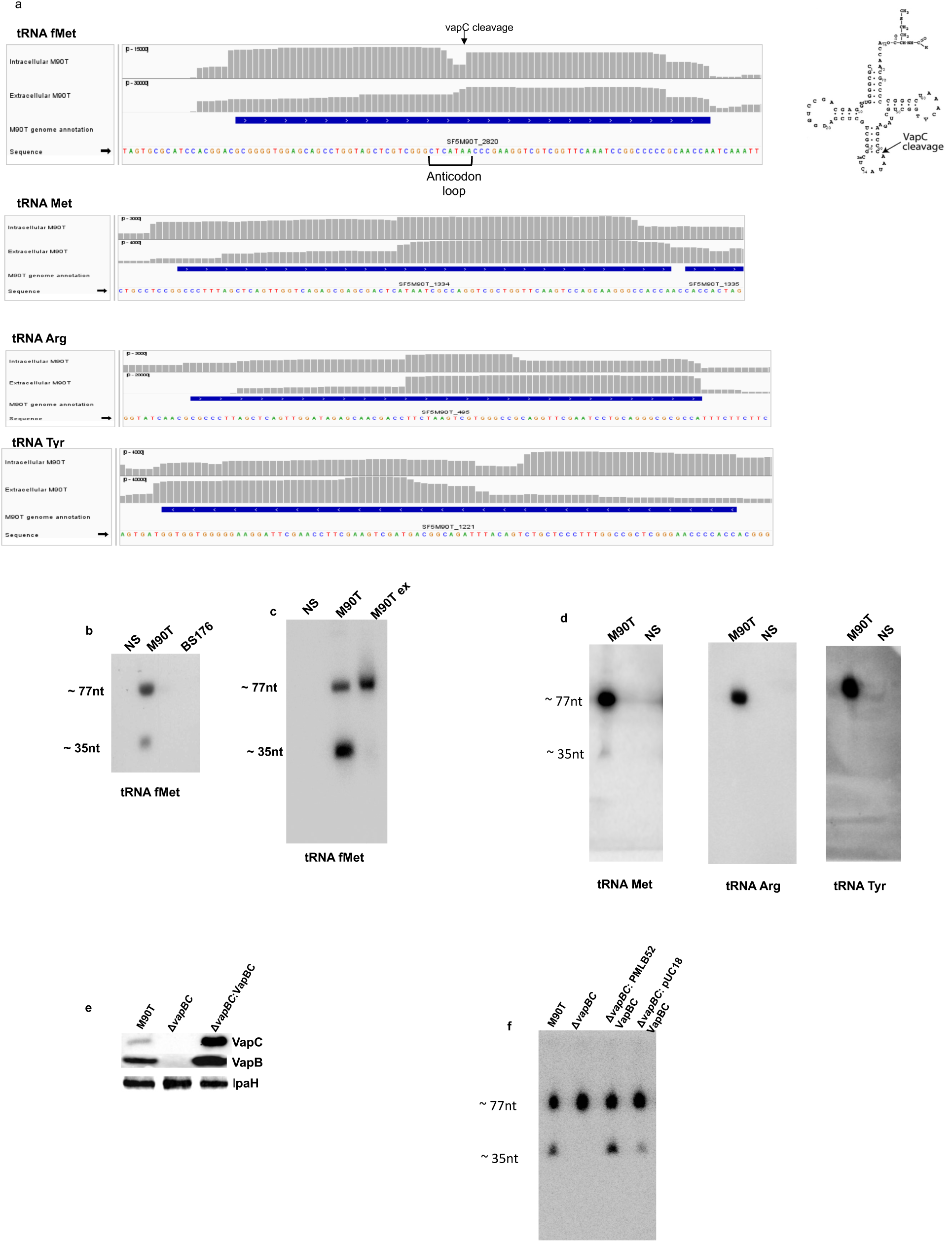
Intracellular Activation of the VapBC Toxin-Antitoxin System in *Shigella flexneri* Leads to Specific Cleavage of tRNA^fMet. **(a) Anticodon Loop Cleavage of tRNA^fMet^ in Intracellular *Shigella flexneri*** RNA-seq coverage plots extracted from dual RNA-sequencing of enterocytic HCT116 cells infected with *Shigella flexneri* M90T for 5 hours, comparing intracellular and extracellular bacterial transcriptomes. Grey bars represent read coverage across individual tRNA genes. Intracellular *S. flexneri* shows a marked drop in read abundance specifically at the anticodon loop region of the initiator tRNA^fMet^ (indicated by black arrows), consistent with VapC-mediated cleavage. No such cleavage pattern is observed in extracellular bacteria or other tRNA, supporting the intracellular-specific activation of VapC ribonuclease activity. Blue arrows indicate annotated tRNA gene orientation and coordinates. **(b-c) Northern Blot Confirmation of tRNA^fMet^ Cleavage During Infection** Northern blot analysis of bacterial tRNA^fMet^ from HCT116 human epithelial cells infected 5 hours with Shigella flexneri. Total RNA was extracted 5 hours post-infection. Cleavage of tRNA^fMet was assessed using a probe targeting the 3’ end of the transcript. Lane annotations: NS – Non-infected HCT116 cells (negative control); M90T – Cells infected with wild-type *S. flexneri* M90T, BS – Cells infected with the noninvasive *S. flexneri* strain BS176, M90T ex – RNA from extracellular *S. flexneri* M90T cultured in cell medium **(d) Northern Blot Confirmation of the specificity of tRNA^fMet^ Cleavage During Infection** Northern blot analysis of the indicated bacterial tRNA from HCT116 human epithelial cells infected 5 hours with *Shigella flexneri.* Membranes were probed for bacterial tRNA Methionine, tRNA Arginine and tRNA Tyrosine. tRNA^Arg and tRNA^Tyr remained intact, in coherence with the selective activity of VapC towards tRNA^fMet^. **(e) Western blot analysis of VapB and VapC expression** Western blot showing the expression of VapB and VapC in *Shigella flexneri* wild-type (WT), deletion mutant (Δ*vapBC*), and complemented (Comp) strains during in vitro growth at the stationary phase. Home-made rabbit polyclonal antibodies were used to detect VapB and VapC. The Δ*vapBC* mutant shows no detectable signal for either protein, confirming both successful gene deletion and antibody specificity. Expression of VapB and VapC is restored in the complemented strain. IpaH was used as a loading control. **(f) Northern blot analysis of tRNA^fMet^ cleavage in *Shigella* shows VapC-dependent activity.** Northern blot detection of tRNA^fMet^ in *Shigella flexneri* strains to assess VapC-dependent cleavage activity. HCT116 epithelial cells were infected for 5 hours with wild-type (WT), Δ*vapBC* mutant, and two complemented strains: Δ*vapBC*::pMLB52VapBC (low-copy plasmid) and Δ*vapBC*::pUC18VapBC (high-copy plasmid). RNA was extracted from intracellular bacteria and analyzed using a probe specific to the 3ʹ region of tRNAf^Met^.

Consistent with the reported VapC activity *in vitro*, cleavage of tRNA*^fMet^* was specific in infected cells, sparing other tRNAs in intracellular bacteria, as shown by both transcriptome and northern blot analyses (**Figures 1a and 1d**). To directly demonstrate the role of the VapBC TA system in this cleavage, a Δ*vapBC* mutant strain and complemented strains (Δ*vapBC*-pUCVapBC, Δ*vapBC*-PMLB52VapBC) were generated (**Figure 1e**). Likewise, infection of HCT116 cells with the indicated strains showed an abrogation of tRNA*^fMet^* cleavage when infected with the Δ*vapBC* mutant restored with the complemented strains (**Figure 1f**). Altogether, these results showed that the VapBC operon is activated upon *Shigella* infection.

### Environmental stress differentially regulates expression and activation of the *Shigella flexneri* VapBC TA

The VapBC toxin–antitoxin (TA) system is encoded by a bicistronic operon that produces an mRNA encoding both the VapB antitoxin and the VapC toxin, as well as an antisense transcript named *trbH* (as-*trbH*), whose function remains unknown (**Figure 2a**). The system is subject to autoregulation: the VapBC protein complex binds to inverted repeats in its own promoter region, typically repressing its own transcription (K. S. Winther & Gerdes, 2012) (**Figure 2a**).

**Figure 2.**
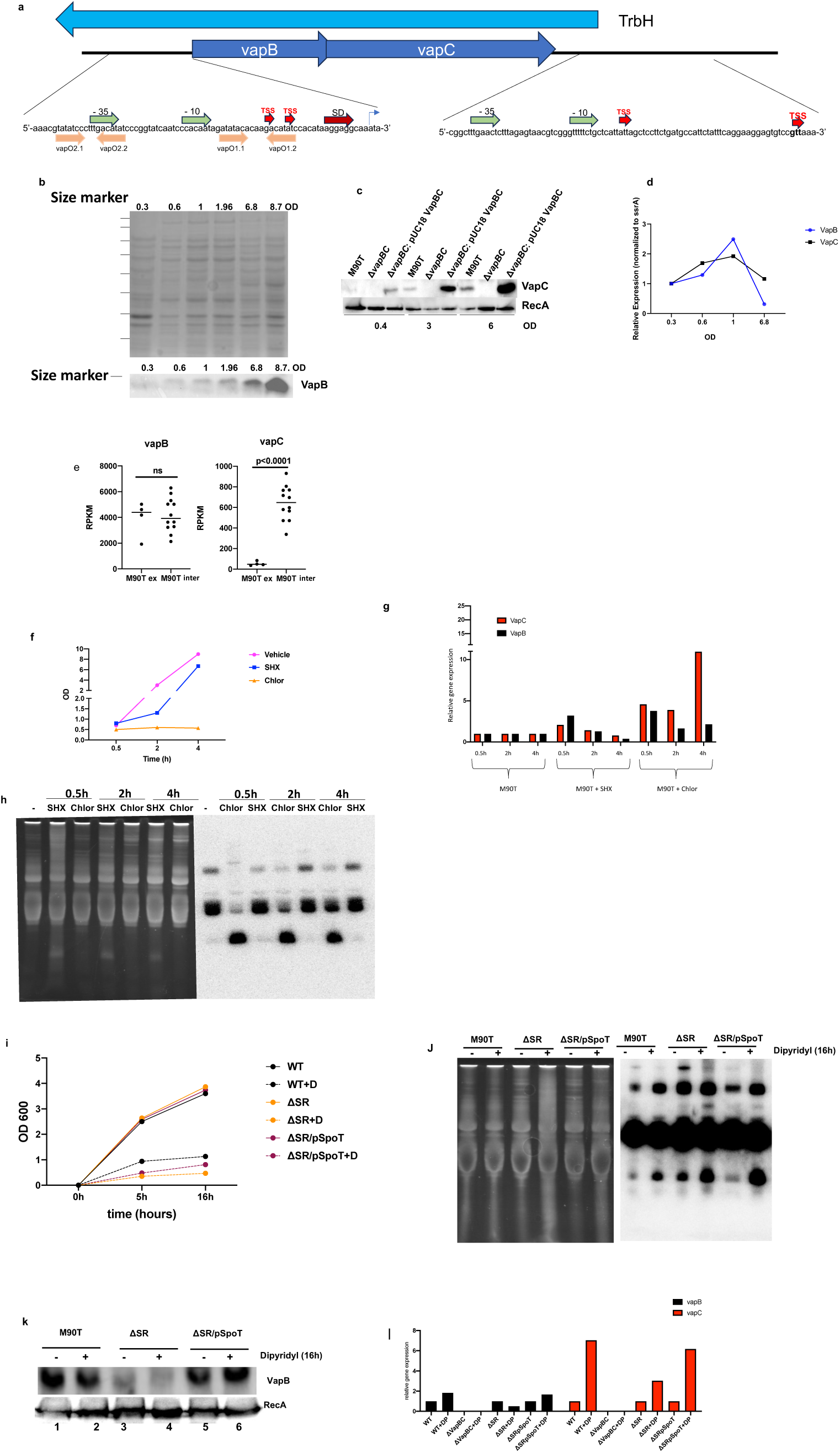
Environmental Cues and Stringent Response Shape *vapBC* Operon Expression and Toxin Activation in *Shigella flexneri*. **(a) Schematic representation of the *vapBC* operon** The positions of the *vapB* and *vapC* genes are indicated by blue arrows, with the adjacent *trbH* gene shown upstream. Below, the nucleotide sequences upstream of *vapBC* are expanded to show predicted promoter elements. The −35 and −10 boxes for each promoter are indicated by green arrows, and transcription start sites (TSS) are marked in red. The Shine-Dalgarno (SD) sequence is shown in blue. Individual promoters (vapO2.1, vapO2.2, vapO1.1, vapO1.2) are indicated beneath their respective sequences **(b–d) Analysis of VapB and VapC Expression Levels According to Bacterial Culture Density** (**b**) Western blot showing VapB protein levels at different optical densities (OD), detected using homemade polyclonal antibodies. Ponceau S staining was used to confirm equal protein loading across samples. **(c)** Western blot of VapC protein levels across the same OD series, detected using homemade antibodies and normalized to RecA as a loading control. **(d)** RT-qPCR analysis of *vapB* and *vapC* transcript levels over time, normalized to the housekeeping RNA *ssrA*. **(e) Differential Expression of *vapB* and *vapC* During Intracellular versus Extracellular Growth of *Shigella*** RNA-seq analysis comparing *vapB* and *vapC* expression levels in *Shigella flexneri* M90T grown intracellularly (M90T int) versus extracellularly (M90T ex). Expression values are presented as RPKM (Reads Per Kilobase per Million mapped reads). Statistical significance was determined using a two-sided Student’s *t*-test; indicated *p*-values reflect differences in expression between conditions. **(f) Growth kinetics of *Shigella* in BTCS medium under SHX and chloramphenicol treatment.** Growth curve of *Shigella flexneri* M90T (WT) cultured in BTCS medium under untreated conditions (vehicle) or treated with SHX (serine hydroxamate 100µg/mL) or chloramphenicol (Chlor 30µg/mL). Bacterial growth was monitored by measuring OD600 over time. **(g) Expression levels of *vapB* and *vapC* under stress conditions induced by SHX and chloramphenicol in *Shigella*** Quantitative RT-PCR analysis of *vapB* and *vapC* expression in *Shigella flexneri* M90T at 0.5, 2, and 4 hours post-treatment under three conditions: untreated control, SHX treatment (100 µg/mL serine hydroxamate), and chloramphenicol treatment (30 µg/mL). Expression levels were normalized to 16S rRNA and are presented as relative fold changes compared to the untreated 0.5-hour time point. **(h) Northern blot tRNAf^Met^ under SHX and chloramphenicol treatments** Total RNA was extracted from *Shigella flexneri* M90T cells treated with SHX (serine hydroxamate 100µg/mL) or chloramphenicol (Chlor 30µg/mL) for 0.5, 2, and 4 hours. Left panel: Ethidium bromide-stained gel showing total RNA prior to transfer, used as a loading control.Right panel: Northern blot probed with a specific oligonucleotide targeting initiator tRNA^fMet^. Untreated cells (indicated as“–”) were used as a reference control. **(i) Bacterial Growth Kinetics of *Shigella* Wild-Type, Δ*spoT*Δ*relA* Mutant, and Complemented Strain Under Dipyridyl Treatment** Growth curves of *Shigella flexneri* strains cultured in LB medium in the absence (vehicle control) or presence of 250 µM 2,2ʹ-dipyridyl (+D), an iron chelator. Optical density at 600 nm (OD₆₀₀) was measured at regular intervals to monitor bacterial growth. The strains tested included the wild-type *S. flexneri* M90T (WT), a double knockout mutant lacking both *spoT* and *relA* (ΔSR), and a complemented strain (ΔSR/pSpoT) carrying a plasmid expressing *SpoT*. **(j) Northern Blot Analysis of tRNA^fMet^ in *Shigella* Strains Under Dipyridyl Treatment** Total RNA was extracted from *Shigella flexneri* strains grown for 16 hours in LB medium with (+) or without (−) 250 µM 2,2ʹ-dipyridyl and analyzed by Northern blot. Left panel: Ethidium bromide-stained gel showing total RNA prior to transfer, used as a loading control. Right panel: Northern blot probed with a specific oligonucleotide targeting initiator tRNA^fMet **(k) Western Blot Analysis of VapB Expression in *Shigella* Strains Under Dipyridyl Treatment** Western blot showing VapB protein levels in *Shigella flexneri* wild-type (WT), Δ*SR* mutant (Δ*spoT* Δ*relA*), and Δ*SR*/pSpoT complemented strains after 16 hours of growth in LB medium, with ( + ) or without (–) 250 µM 2,2ʹ-dipyridyl. RecA was used as a loading control. **(l) Expression Levels of *vapB* and *vapC* in *Shigella* Strains Under Dipyridyl Treatment** Quantitative RT-PCR analysis of *vapB* and *vapC* transcript levels in *Shigella flexneri* wild-type (WT), Δ*SR* mutant (Δ*spoT* Δ*relA*), and *ΔSR*/pSpoT complemented strains after 16 hours of growth in LB medium, in the absence or presence (+DP) of 250 µM dipyridyl. Gene expression levels were normalized to 16S rRNA and are presented as relative fold change compared to untreated WT

Consistently, *vapBC* transcript levels decrease as VapB and VapC protein levels rise with increasing culture density, as shown by RT-qPCR and immunoblot analyses, respectively (**Figure 2b–d**). In the transcriptome of intracellular *Shigella*, we observed increased expression of *vapC* mRNA compared to extra-cellular *Shigella*, while *vapB* mRNA expression levels remained unchanged (**Figure 2e**). This suggests that the functional activation of the operon is accompanied by transcriptional induction of the toxin, while antitoxin levels remained barely affected.

To better understand this regulatory coupling, we further investigated how various stressors affect the transcription and activity of the VapBC operon. We thus subjected *Shigella* to two well-studied stresses, the stringent response following amino acid starvation, which can be rapidly induced via addition of serine hydroxamate (SHX), and translation inhibition, induced by treatment with chloramphenicol. Chloramphenicol treatment exerts severe growth inhibition and leads to a rapid upregulation of *vapBC* transcripts **(Figures 2f-g)**. Notably, while *vapC* mRNA expression level remained elevated throughout the stimulation period, *vapB* mRNA levels progressively declined **(Figure 2g).** In parallel, northern blot analysis revealed rapid cleavage of tRNA*^fMet^* over-time, indicating activation of VapC toxin activity in response to chloramphenicol treatment **(Figure 2h**). Stimulation by SHX led to a transient growth inhibition, more likely linked to its reported biotic degradation (Patacq et al., 2020) (**Figure 2f**). Likewise, SHX stimulation triggered a rapid but transient expression of *vapBC* mRNAs, albeit less sustained than that observed with chloramphenicol treatment **(Figure 2g).** This transcriptional induction was not associed to tRNA*^fMet^* cleavage, as shown by northern blot (**Figure 2h**).

Since SHX-induced amino acid primarily activates the RelA-mediated stringent response, we investigated whether the SpoT-mediated stringent response, recently implicated in *Shigella* virulence (Kago et al., 2023), could also activate the *vapBC* operon. To this end, we tested whether the iron-chelating agent dipyridyl, known to activate SpoT, could trigger tRNA*^fMet^* cleavage, and whether this effect was dependent on SpoT. Wild-type bacteria, the corresponding Δ*relA* Δ*spoT* (*ΔRS*) mutant, and the ΔRS strain reconstituted with the wild-type *spoT* gene (pWKS30-*spoT*) were subjected to iron limitation using dipyridyl. Interestingly, northern blot revealed that the *ΔRS* mutant exhibited increased basal tRNA*^fMet^* cleavage (**Figure 2j, compare lane 3 to lane 1**). Importantly, this strain also showed a marked reduction in VapB protein levels (**Figure 2k, lanes 3-4**), while *vapB* mRNA levels remained unaffected (**Figure 2l**), indicative of a post-transcriptional mecanism for the defect in VapB expression in the absence of SpoT. Dipyridyl stimulation resulted in growth inhibition for the 3 strains and tRNA*^fMet^* cleavage, indicating activation of the *vapBC* operon under iron-limiting conditions (**Figures 2i-j**). Despite operon activation, VapB protein levels remained relatively stable in SpoT-proficient backgrounds (i.e., wild-type and complemented *ΔRS*) (**Figure 2k, compare lanes 1-2 and lane 5-6**), while VapB protein level remained hardly detectable in the *ΔRS* mutant (**Figure 2k compare lanes 3 to 4**).

While dipyridyl did not induce major change in *vapB* transcript levels, it led to a strong increase in *vapC* mRNA in all strains, with enhanced expression in SpoT-proficient backgrounds (i.e., wild-type and complemented ΔRS) (**Figure 2l**).

Altogether, these findings demonstrate that the *Shigella flexneri* VapBC system is differentially regulated by environmental stress, and that transcriptional induction is not always coupled with VapC activation. The stringent response, primarily mediated by SpoT, appears to influence operon responsiveness by maintaining VapB antitoxin expression levels. Iron limitation emerges as a strong activator of the *vapBC* operon and is associated with a pronounced upregulation of *vapC* mRNA.

### VapBC Promotes Intracellular Activation of the Type III Secretion System

To investigate whether VapBC plays a role in *Shigella flexneri* virulence, we examined the impact of *vapBC* deletion on T3SS activation. First, we verified that deletion of *vapBC* did not affect the stability of the virulence plasmid pINV, either in the inoculum (i.e., the bacterial population used for infection) or 16 hours post-infection in HCT116 cells. A colony assay on Congo red agar plates showed no difference in pINV stability, with both the wild-type and Δ*vapBC* mutant forming red colonies, suggesting intact plasmid retention both prior to and during infection (**Supplementary Figure 1a-c**). The T3SS machinery also appeared functional *in vitro*, as Congo red-induced secretion assays revealed comparable release of early (IpaC) and late (IpaH) effectors between two strains **(Supplementary Figure 1d-e**).

To assess the impact of VapBC on T3SS activity in intracellular bacteria, we combined dual RNA-seq transcriptomic profiling with a single-cell approach using the TSAR system to monitor T3SS activity (Campbell-Valois et al., 2014), and Galectin-3-eGFP to track the internalization process and vacuolar escape (Y. Y. Chang et al., 2020).

RNA-seq data showed no major differences in pINV gene expression in the *ΔvapBC* strain (**Table S2 and Figure 3a**), and quantification of the time-lapse images indicated that WT and the *ΔvapBC* strain reached the cytosol at the same time (**Figure 3f-g**). However, we noted a delay in the unpeeling of BCV membranes upon initial vacuolar rupture in a subset of entering mutant bacteria, that remains closely connected with the broken BCV remnants. At later stage, reporter-based flow cytometry revealed a consistent reduction in the number of intracellular bacteria activating their T3SS at 5 hours post-infection (**Figure 3d-e**). This defect was further corroborated by plaque assays, which demonstrated a drastic decrease in plaque formation in the absence of VapBC, despite normal invasion efficiency (data not shown). Together, these data indicate that VapBC is required for efficient intracellular T3SS activation.

**Figure 3.**
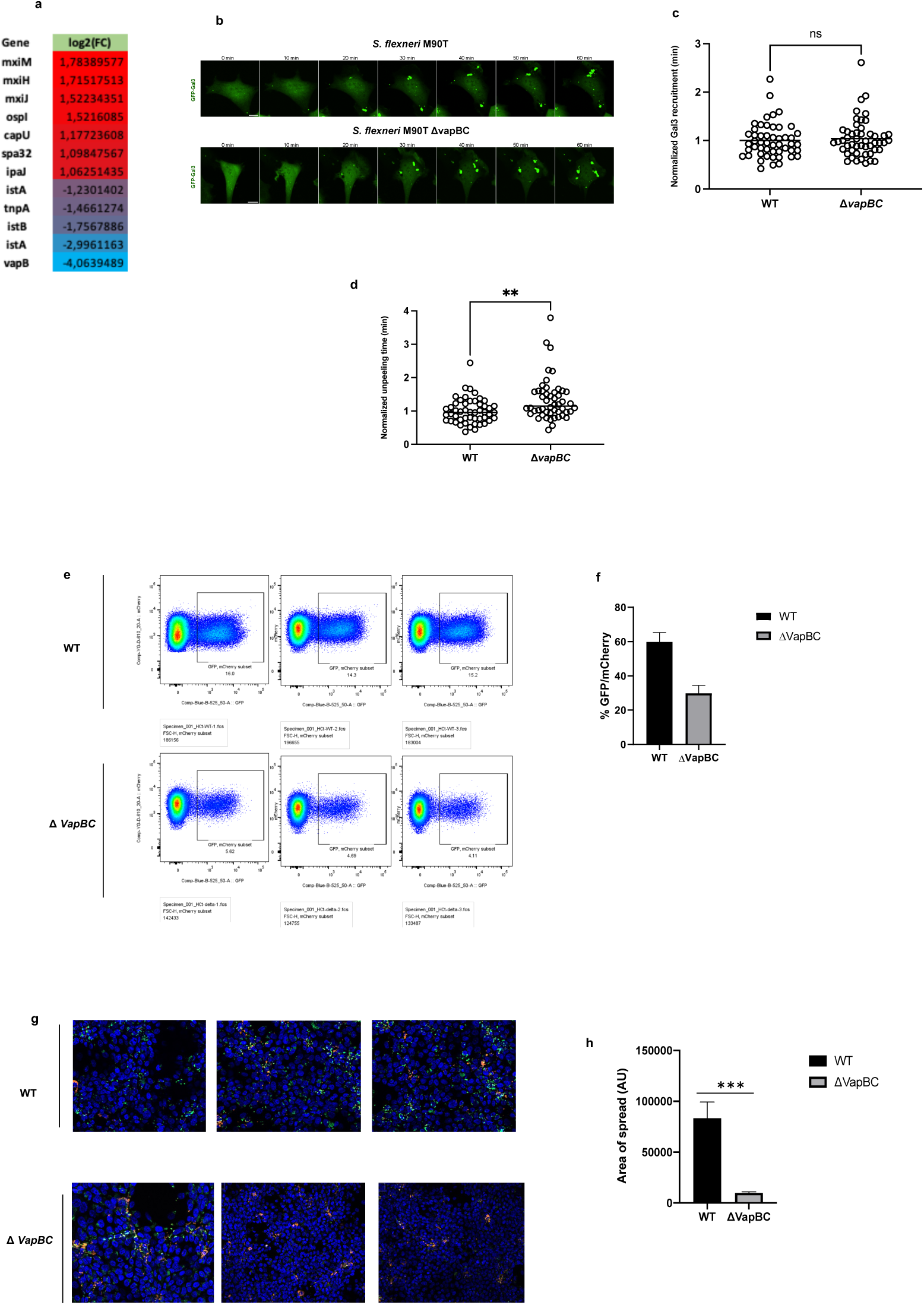
VapBC Supports T3SS Activation and Intracellular Virulence in *Shigella flexneri*. **(a) Differential expression of virulence plasmid-encoded genes in *Shigella flexneri* during infection of HCT116 cells.** Heatmap showing log₂ fold change (log₂(FC)) of selected virulence plasmid genes significantly differentially expressed between wild-type (WT) and *ΔvapBC Shigella flexneri* strains at 5 hours post-infection of HCT116 epithelial cells, based on dual RNA-seq analysis. Differential expression analysis was performed using DESeq2, and only genes with adjusted p-value < 0.05 are shown. **(b) Live-cell imaging of galectin-3 recruitment during infection of HeLa cells with *Shigella flexneri* WT or *ΔvapBC* strains**. Time-lapse fluorescence microscopy of HeLa cells stably expressing galectin-3-eGFP infected with *Shigella flexneri* wild-type (WT, dsRed) or ΔvapBC mutant (dsRed). Images were acquired every 2 minutes starting from the initial contact of the bacteria with host cells for 120 minutes, and representative frames were merged for each condition. **(c-d) Quantitative Analysis of Galectin-3 Recruitment Dynamics and Vacuolar Unpeeling in Cells Infected with *Shigella flexneri* WT or Δ*vapBC* Strains** Galectin-3 recruitment was used as a marker to quantify the timing of vacuolar rupture (peak Gal3 signal) and the onset of unpeeling (defined as the morphological separation of the Gal3 signal from the entering bacteria). Image analysis was performed using Fiji (ImageJ). Statistical comparisons were made using a two-tailed Student’s *t*-test. p < 0.01; ns, not significant. **(e–f) Flow Cytometry Analysis of Bacterial Populations Expressing the Dual-Fluorescent T3SS Reporter TSAR3.1** The TSAR system uses mCherry expression under the constitutive *r16s* ribosomal promoter to mark bacteria, and GFP expression under the *ipaH* promoter as a readout of Type III Secretion System (T3SS) activation. HCT116 epithelial cells were infected for 5 hours at an MOI of 50 with *Shigella* strains harboring the TSAR3.1 reporter. Intracellular bacteria were recovered and analyzed by flow cytometry. **(e)** Representative flow cytometry plots showing GFP (T3SS activity) versus mCherry (bacterial marker) fluorescence for wild-type (WT) and Δ*vapBC* strains across three biological replicates. **(f)** Quantification of the percentage of double-positive (GFP⁺/mCherry⁺) bacteria, reflecting the proportion of the population with active T3SS. **(g–h) Immunofluorescence Microscopy and Quantification of Bacterial Spread** **(g)** HCT116 cells were infected for 5 hours with TSAR-expressing *Shigella* strains. mCherry (red) marks total bacteria, while GFP (green) indicates T3SS activation. Host cell nuclei were stained with DAPI (blue). **(h)** Bar graph quantifying the bacterial spread area (arbitrary units, AU) for WT and Δ*vapBC* strains. Image analysis was performed using Fiji (ImageJ). Statistical significance was assessed using a two-tailed Student’s t-test. ***p < 0.001.

### VapBC controls the expression of key genes Involved in metabolism and stress responses

We further investigated whether VapBC plays a role in *Shigella’s* adaptation to its intracellular lifestyle. While inactivation of *vapBC* did not affect bacterial growth in standard microbiological media, the intracellular growth of the *ΔvapBC* strain was impaired, as determined by colony-forming unit (CFU) counts post-infection (**Figure 4a**). We hypothesized that this intracellular growth defect could result from altered expression of genes involved in metabolism, stress responses, or other pathways essential for bacterial adaptation within host cells.

**Figure 4.**
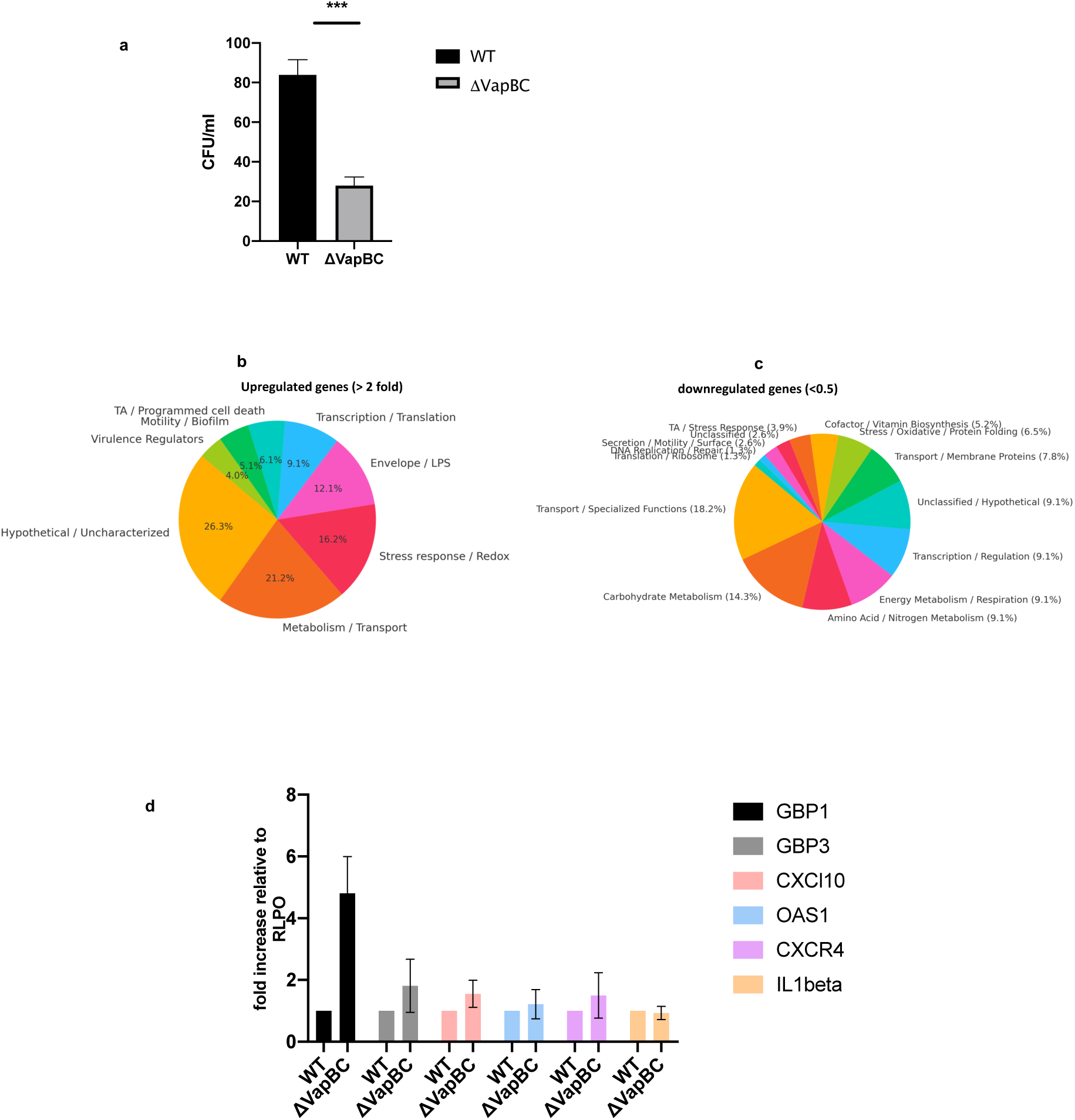
VapBC Supports Intracellular Adaptation of Shigella Through Bacterial and Host Transcriptional Modulation. **(a) Intracellular Survival of WT and Δ*vapBC Shigella* in HCT116 Cells** Colony-forming unit (CFU) assay quantifying intracellular bacteria recovered from HCT116 cells 16 hours post-infection at a multiplicity of infection (MOI) of 0.5 (n = 3). The wild-type (WT) strain exhibits significantly higher intracellular survival compared to the Δ*vapBC* mutant, Statistical significance was assessed using a two-tailed Student’s t-test. ***p < 0.001. **(b-c) Functional classification of chromosomally encoded genes differentially expressed in Δ*vapBC* versus WT *Shigella flexneri* during intracellular infection.** Pie charts represent the functional categories of chromosomal genes significantly differentially expressed in HCT116 cells infected for 16 hours with either wild-type (WT) or Δ*vapBC* strains, based on dual RNA-seq analysis. In (**b**), upregulated genes are shown; in (**c**), downregulated genes are depicted. Genes selected exhibit at least a 2-fold change in expression (log₂ fold change ≥ 1 or ≤ –1) and an adjusted p-value < 0.05, as determined by DESeq2. Functional categories were assigned based on curated gene ontology annotations. **(d) qPCR Analysis of Host Immune Gene Expression in Response to WT and *ΔvapBC Shigella* Infection** Quantitative PCR (qPCR) analysis of immune-related gene expression in HCT116 cells infected for 16 hours with wild-type (WT) or Δ*vapBC Shigella* strains at a multiplicity of infection (MOI) of 0.5. Total RNA was extracted and analyzed for expression of GBP1, GBP3, CXCL10, OAS1, CXCR4, and IL1β, with normalization to RPLP0 as a housekeeping gene. Data are presented as fold change relative to RPLP0 expression. Error bars indicate standard deviation. Data are representation of n=2 experiments.

To explore the basis of this phenotype, we analyzed transcriptional differences between the wild-type and Δ*vapBC* strains during infection **(Figure 4b-c).** Transcriptomic analysis of intracellular Δ*vapBC* mutant revealed strong downregulation of genes essential for core metabolic processes. Among the most significantly repressed genes was *thiC*, involved in thiamine (vitamin B1) biosynthesis, a vital cofactor for carbohydrate metabolism. *ubiA*, which encodes a key enzyme in the biosynthesis of ubiquinone (coenzyme Q), was also affected, suggesting impaired electron transport and energy production.

Several additional repressed genes further highlight defects in biosynthetic and cellular maintenance functions. *glyQ* (glycyl-tRNA synthetase α-subunit) and *cysS* (cysteinyl-tRNA synthetase), both critical for accurate tRNA charging and protein synthesis, were suppressed, indicating potential translational stress. Genes required for amino acid biosynthesis and transport, such as *asnC*, *cysP*, *cysS*, *hisP*, and *leuC*, were repressed, potentially limiting the availability of key building blocks for protein synthesis. In parallel, the downregulation of *asnC*, a regulator of asparagine metabolism, and *ddpX*, a D-Ala-D-Ala dipeptidase, points to disrupted amino acid and peptide utilization. Moreover, *iucB*, a component of the aerobactin siderophore biosynthesis system, was also repressed, suggesting impaired iron acquisition, a known bottleneck for *Shigella* during intracellular survival.

Transcriptomic analysis of the Δ*vapBC* mutant also revealed the upregulation of a large set of genes associated with stress response, membrane composition, and TA systemes adaptation Genes involved in LPS modification and envelope biosynthesis, such as *arnF*, *waaO*, *wecH*, *wzy*, *wzzB*, and *rfbG*, were significantly induced, suggesting altered outer membrane architecture. This was accompanied by increased expression of *ompC*, a major outer membrane porin, and *murC*, involved in peptidoglycan biosynthesis, pointing to cell envelope remodeling. A number of oxidative and redox stress-associated genes were also upregulated, including *dsbC*, *oxyR*, *sufA*, *sufE*, *sapB*, *uspF*, *hslV*, and *gadX*, consistent with activation of defense mechanisms against intracellular stress. Similarly, *nhaA* and *nudB*, involved in ion and pH homeostasis, were elevated. We also observed upregulation of type I and II TA module components, including *higA*, *hokD*, *ortT*, *yafO*, *yjbD*, and *ymiC*, possibly highlighting compensatory activation of alternative TA systems in the absence of VapBC.

Overall, loss of VapBC induces a transcriptomic profile marked by metabolic repression and activation of stress-response pathways, a signature broadly associated with reduced bacterial fitness under stress conditions.

### VapBC contributes to bacterial fitness by restricting cell-autonomous Immunity

To further acess the contribution of *vapBC* in the fate of the intracellular bacterium, we analyzed the host transcriptome response. Transcriptomic profiling of host cells infected with the Δ*vapBC* mutant strain of *Shigella flexneri* revealed a marked upregulation of numerous interferon-stimulated genes (ISGs) compared to the wild-type strain. This response encompassed genes involved in innate immune sensing (*IFIH1*, *DHX58*, *DDX60*), interferon signaling and regulation (*IRF1*, *IRF7*, *IRF9*), and antiviral effectors (*ISG15*, *OASL*, *IFIT1–3*, *RSAD2*, *MX1/2*). In addition, we observed increased expression of chemokines and cytokines such as *CXCL10*, *CXCL11*, and *IL15*, as well as immune regulators (*TRIM21*, *TRIM25*, *TRIM5*, *CASP1*), all consistent with a robust type I and type II interferon response (**Table S3**). Importantly, guanylate-binding proteins (GBPs), which belong to a large IFN-induced GTPase family, and known to restrict *Shigella* dissemination (Wandel et al., 2017) were at the top ten gene upregulated in the vapBC mutant, was further confirmed by RT-qPCR (**Figure 4d**). Collectively, these findings demonstrate that VapBC expression is important for *Shigella flexneri*’s host immune responses evasion during infection.

## Discussion

Our study uncovers a central role for the VapBC operon in the intracellular adaptation of *Shigella flexneri*. We demonstrate that the operon is specifically activated during infection of host epithelial cells, as clearly demonstrated by the detection of VapC-dependent ribonuclease activity targeting the initiator tRNA*^fMet^*, an effect abolished in the *ΔvapBC* mutant, confirming VapC as the active effector.

Inside host cells, *Shigella flexneri* encounters a highly restrictive environment, limited in key nutrients such as amino acids, sugars, and iron. In response, the bacterium undergoes a major metabolic reprogramming, shifting from oxygen-dependent respiration to mixed-acid fermentation, which helps support intracellular survival and the expression of virulence factors (Lucchini et al., 2005)(Pieper et al., 2013)(Kentner et al., 2014; Koestler et al., 2018). These nutrient limitations, particularly amino acid and iron deprivation, are classical triggers of the bacterial stringent response alarmone (p)ppGpp (W. Li et al., 2015; Vinella et al., 2005).

Recent reports have highlighted a central role for the stringent response, particularly through the RSH protein SpoT, in promoting the expression of the key *Shigella* virulence regulator VirF and its downstream genes. SpoT is a bifunctional (p)ppGpp synthase and hydrolase responsible for maintaining basal levels of (p)ppGpp (Fernández-Coll & Cashel, 2020; Laffler & Gallant, 1974). In the absence of SpoT, we observed a marked reduction in VapB protein levels despite unchanged *vapB* mRNA expression, strongly suggesting that SpoT actively promotes VapB accumulation, likely through post-transcriptional mechanisms (e.g., enhancing mRNA stability or translation efficiency) or post-translational regulation (e.g., preventing protein degradation). This imbalance may result in unrestrained VapC activity, as indicated by increased basal tRNA*^fMet^* cleavage in the Δ*relA* Δ*spoT* strain. These findings suggest a role for SpoT in maintaining TA system balance under basal conditions, possibly acting as a safeguard against inappropriate toxin activation.

SpoT is known to be activated by deficiencies in carbon and iron, two conditions encountered by *Shigella flexneri* during intracellular infection (Vinella et al., 2005)(Meyer et al., 2021). In this study, we identified iron starvation as a strong activator of *vapBC* operon activity, pointing to a plausible, though likely not exclusive, mechanism for *in vivo* operon activation. This activation was associated with a pronounced transcriptional upregulation of *vapC*, without corresponding changes in *vapB* transcript levels or VapB protein abundance. This selective upregulation was unexpected, as TA systems are typically organized as bicistronic operons producing a single mRNA encoding both the antitoxin and the toxin.

This observation suggests that stress-induced activation of the stringent response via SpoT, particularly under iron-limiting conditions, may decouple *vapC* and *vapB* expression. Similar imbalances have been described in type I TA systems, where stress conditions favor increased toxin stability or translation. For example, the *aapA1* toxin mRNA of *Helicobacter pylori* is more stable than its cognate antitoxin under oxidative stress (Mortaji et al., 2020) and in *E. coli*, editing of *hokB* mRNA increases with cell density, resulting in more active toxin isoforms (Bar-Yaacov et al., 2017; Wilmaerts et al., 2019). By analogy, the disproportionate upregulation of *vapC*, in the absence of a parallel increase in *vapB*, likely disrupts the stoichiometric balance between toxin and antitoxin. This could lead to the accumulation of free VapC toxin, as basal levels of VapB may be insufficient to neutralize the excess VapC, thereby enabling ribonuclease activity upon stress-induced iron limitation.

In challenging environments, such as host infection, metabolic adjustments are essential for microorganism survival and competitiveness. We showed that intracellular growth was affected upon *vapBC* inactivation, and the transcriptional landscape reveals a broad repression of metabolic pathways. Notably, genes associated with amino acid biosynthesis, carbohydrate utilization, energy production, and cofactor synthesis were consistently repressed. This transcriptional shift may compromise the bacterium’s ability to synthesize proteins, maintain redox balance, and produce ATP efficiently, all essential functions for replication and T3SS activation within host cells. Notably, T3SS assembly and effector secretion are energetically demanding processes tightly linked to central metabolism, as shown in *Pseudomonas aeruginosa* (Gil-Gil et al., 2023).

Consistently, we observed that bacterial virulence was primarily affected at later stages of infection, particularly during intercellular dissemination, while the invasion step remained globally unaffected. While the transcriptome of intracellular bacteria did not reveal major changes in the expression of genes encoded by the virulence plasmid, this was accompanied by a reduced proportion of intracellular bacteria activating their T3SS, indicating that VapBC contributes to sustaining T3SS activation during intracellular spreading.

In Shigellosis, the injection of late effectors by the T3SS is essential for evasion of host immune defenses, particularly mechanisms of cell-autonomous immunity (López-Jiménez et al., 2024). We show that in the absence of *vapBC*, *Shigella* is more readily detected by the host immune system, as indicated by a potent upregulation of interferon-responsive genes. Among these, guanylate-binding proteins (GBPs) are strongly induced. GBPs recognize cytosolic bacteria by binding directly to lipopolysaccharide (LPS), encapsulate the pathogen, promote membrane disruption, and trigger non-canonical inflammasome activation via caspase-4/11 (Dickinson et al., 2023; Kutsch et al., 2020).

In the context of *Shigella*, GBPs also restrict bacterial dissemination, and T3SS activation is crucial for counteracting GBP-mediated immunity. This immune evasion is mediated by the effector IpaH9.8, an E3 ubiquitin ligase that ubiquitinates GBPs and targets them for proteasomal degradation (Wandel et al., 2017; P. Li et al., 2017). In our study, the robust upregulation of GBPs in Δ*vapBC*-infected cells suggests that VapBC contributes, directly or indirectly, to maintaining effective evasion of GBP-mediated defenses. This may occur by supporting proper T3SS effector delivery or by limiting exposure of pathogen-associated molecular patterns (PAMPs).

Indeed, transcriptomic analysis reveals that in the absence of VapBC, *Shigella* activates a stress-associated transcriptional program aimed at reinforcing the bacterial envelope, possibly as a protective mechanism against envelope-targeting host defense. The upregulation of genes involved in LPS core and O-antigen biosynthesis suggests a compensatory effort to rebuild or stabilize the outer membrane, potentially restoring the protective function of the O-antigen but also immunostimulatory motifs such as lipid A. Concurrent upregulation of outer membrane porins and cell wall biosynthesis genes further supports the idea of envelope remodeling in response to immune pressure.

These adaptations may help the bacterium resist host mechanisms such as guanylate-binding proteins, which target damaged or roughened LPS, and non-canonical inflammasome pathways that sense cytosolic LPS (Kutsch et al., 2020). Altogether, these responses indicate that VapBC may contribute to maintaining envelope integrity under intracellular conditions, and that its absence triggers a broad transcriptional response aimed at limiting membrane damage and immune recognition within the host cytosol. Additionally, the detection of a subset of entering mutant bacteria that remains connected with the broken BCV remnants may also compromise immune evasion, as these structures can serve as signals for host immune damage sensors such as galectins (Mellouk et al., 2024).

While traditionally associated with plasmid maintenance and phage defense, TA systems are increasingly recognized for their involvement in stress adaptation and metabolic regulation (Pizzolato-Cezar et al., 2023). However, their broader physiological relevance remains debated, partly due to studies relying on overexpression models that may not reflect natural regulatory dynamics. Furthermore, many investigations infer TA activation solely from operon transcription, without direct evidence of toxin activity (LeRoux et al., 2020), and our findings emphasize that transcriptional induction of the vapBC operon is not a *proxy* of toxin activity, highlighting the need for direct evidence of molecular function.

In *Shigella* and *Salmonella*, overexpression of VapC toxins leads to global inhibition of translation by specifically cleaving initiator tRNA*^fMet^* (K. S. Winther & Gerdes, 2009, 2011). Recent studies, however, suggest that TA systems, when expressed at native levels, do not necessarily induce global translational arrest or stasis, but instead promote selective remodeling of the bacterial proteome. In *Mycobacterium tuberculosis*, for example, VapC4 cleaves the single tRNA^Cys^ in response to oxidative and copper stress, mimicking cysteine starvation and activating genes involved in cysteine biosynthesis (Barth et al., 2021). This metabolic reprogramming enhances the bacterium’s ability to counter host-derived stress. Similar findings have been reported for the MazF toxin in *M. smegmatis* and *E. coli*, where selective cleavage of RNA leads to fine-tuning of protein synthesis rather than its complete inhibition (Barth & Woychik, 2020; Bezrukov et al., 2021; Nigam et al., 2020; Nikolic et al., 2022). Interestingly, enteric VapC has been proposed to enable translation from non-canonical start codons, potentially favoring the production of stress-adapted proteins (K. S. Winther & Gerdes, 2011) However, this model was based on a synthetic reporter system, and whether VapC activation generates a sufficiently functional proteome *in vivo* remains an open question.

In conclusion, our study reveals a critical role for the *vapBC* operon in the intracellular lifestyle of *Shigella flexneri*. We propose that VapBC contributes to bacterial fitness by enabling transcriptional reprogramming that supports metabolic function, virulence, and resistance to cell-autonomous immunity. The stress-induced transcriptional changes observed upon operon deficiency, including altered expression of envelope and metabolic genes, are consistent with a protective role for VapBC in maintaining intracellular bacterial homeostasis. Further studies are warranted to elucidate the impact of VapBC activity on the *Shigella* intracellular proteome and metabolome, which could reveal specific advantages conferred by this toxin-antitoxin system for bacterial adaptation and persistence within the host.

**Supplementary Figure 1.**
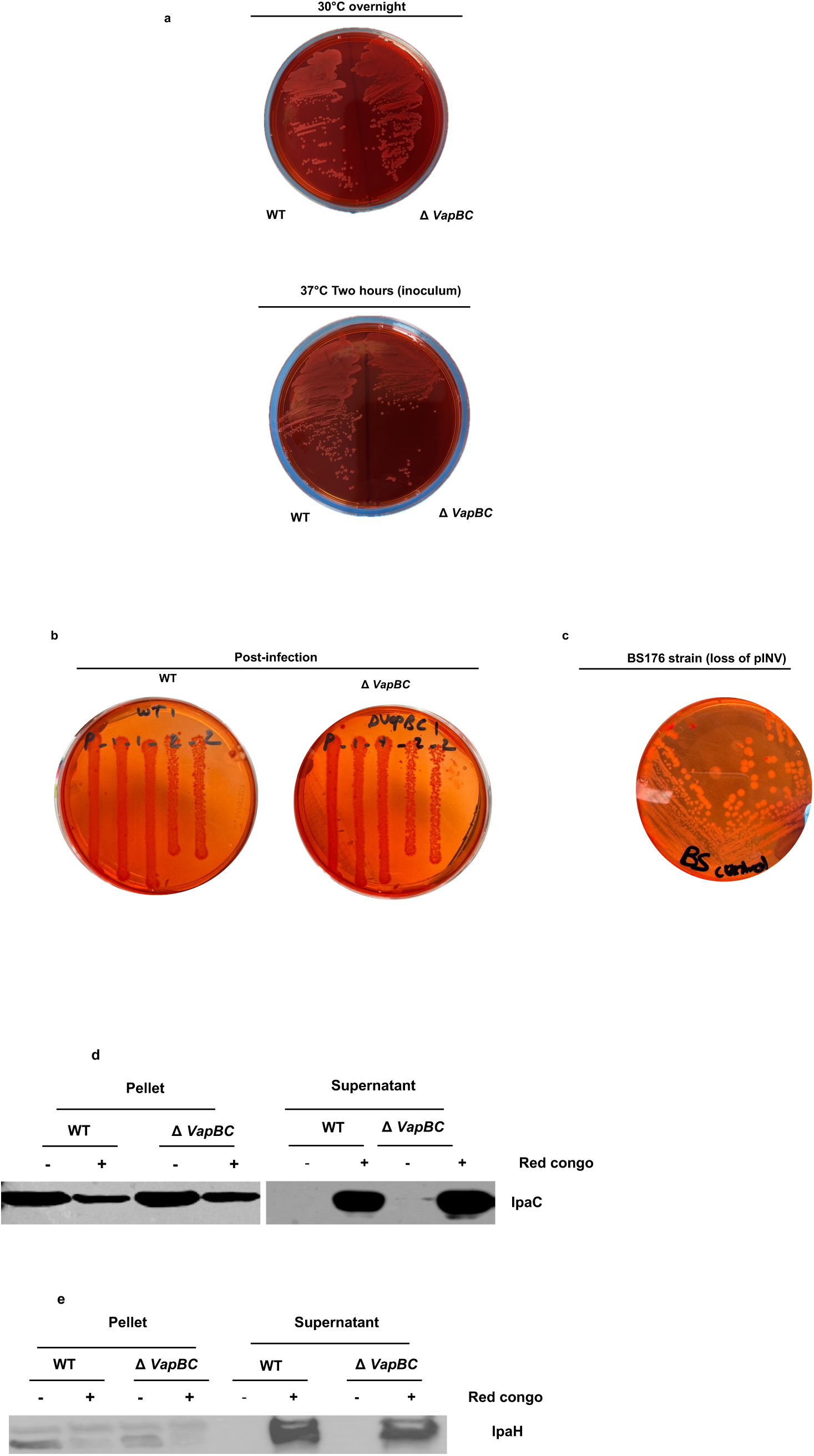
Congo red binding assays in WT and *ΔvapBC Shigella flexneri* strains. **(a)** Top panel: WT and *ΔvapBC Shigella* strains plated side-by-side after overnight incubation at 30 °C. Bottom panel: WT and *ΔvapBC* strains plated side-by-side after 2 hours of incubation at 37 °C, to the bacterial inoculum used for host cell infection. **(b)** Plating of WT and *ΔvapBC* Strains Recovered from Infected Cells on Congo Red Agar WT and ΔvapBC *Shigella* strains were recovered from HCT116 cells 16 hours post-infection and directly plated on Congo red agar to assess virulence plasmid retention. **(c)** Congo Red Binding Control Using Avirulent BS176 Strain : The BS176 avirulent *Shigella* strain, which lacks the virulence plasmid encoding the T3SS, was plated after incubation at 37 °C to serve as a negative control for Congo red binding and colony morphology. **(d)** IpaC expression in the presence (+) or absence (–) of Congo red in various strains. **(e)** IpaH expression in the presence (+) or absence (–) of Congo red in various strains.

**Table S1.**

| NAME | GS<br> follow link to MSigDB | GS DETAILS | SIZE | ES | NES | NOM p-val | FDR q-val | FWER p-val | RANK AT MAX |
| --- | --- | --- | --- | --- | --- | --- | --- | --- | --- |
| HALLMARK_TNFA_SIGNALING_VIA_NFKB | HALLMARK_TNFA_SIGNALING_VIA_NFKB | Details ... | 166 | 0.5362439 | 2.5377135 | 0.0 | 0.0 | 0.0 | 2713 |
| HALLMARK_UV_RESPONSE_DN | HALLMARK_UV_RESPONSE_DN | Details ... | 123 | 0.551196 | 2.4396174 | 0.0 | 0.0 | 0.0 | 3416 |
| HALLMARK_MITOTIC_SPINDLE | HALLMARK_MITOTIC_SPINDLE | Details ... | 197 | 0.46414348 | 2.1316602 | 0.0 | 0.0 | 0.0 | 3136 |
| HALLMARK_TGF_BETA_SIGNALING | HALLMARK_TGF_BETA_SIGNALING | Details ... | 48 | 0.56012034 | 2.044313 | 0.0 | 0.0 | 0.0 | 2730 |
| HALLMARK_INFLAMMATORY_RESPONSE | HALLMARK_INFLAMMATORY_RESPONSE | Details ... | 101 | 0.48493257 | 2.036571 | 0.0 | 0.0 | 0.0 | 3921 |
| HALLMARK_APICAL_JUNCTION | HALLMARK_APICAL_JUNCTION | Details ... | 142 | 0.4190202 | 1.7924135 | 0.0 | 0.0 | 0.0 | 2290 |
| HALLMARK_HYPOXIA | HALLMARK_HYPOXIA | Details ... | 148 | 0.40920293 | 1.7572348 | 0.0 | 0.0 | 0.0 | 2769 |
| HALLMARK_APICAL_SURFACE | HALLMARK_APICAL_SURFACE | Details ... | 25 | 0.50760925 | 1.697089 | 0.0 | 0.0 | 0.0 | 3640 |
| HALLMARK_IL2_STAT5_SIGNALING | HALLMARK_IL2_STAT5_SIGNALING | Details ... | 132 | 0.37687668 | 1.6535618 | 0.0 | 0.0 | 0.0 | 2620 |
| HALLMARK_ESTROGEN_RESPONSE_EARLY | HALLMARK_ESTROGEN_RESPONSE_EARLY | Details ... | 166 | 0.34639362 | 1.6194518 | 0.0 | 0.0 | 0.0 | 3173 |
| HALLMARK_APOPTOSIS | HALLMARK_APOPTOSIS | Details ... | 118 | 0.38299337 | 1.5963557 | 0.0 | 0.0 | 0.0 | 2772 |
| HALLMARK_EPITHELIAL_MESENCHYMAL_TRANSITION | HALLMARK_EPITHELIAL_MESENCHYMAL_TRANSITION | Details ... | 113 | 0.3478472 | 1.5098261 | 0.0 | 0.008578431 | 0.1 | 3267 |
| HALLMARK_KRAS_SIGNALING_UP | HALLMARK_KRAS_SIGNALING_UP | Details ... | 111 | 0.41266316 | 1.4708889 | 0.0 | 0.014485155 | 0.2 | 2970 |

**Table S2.**
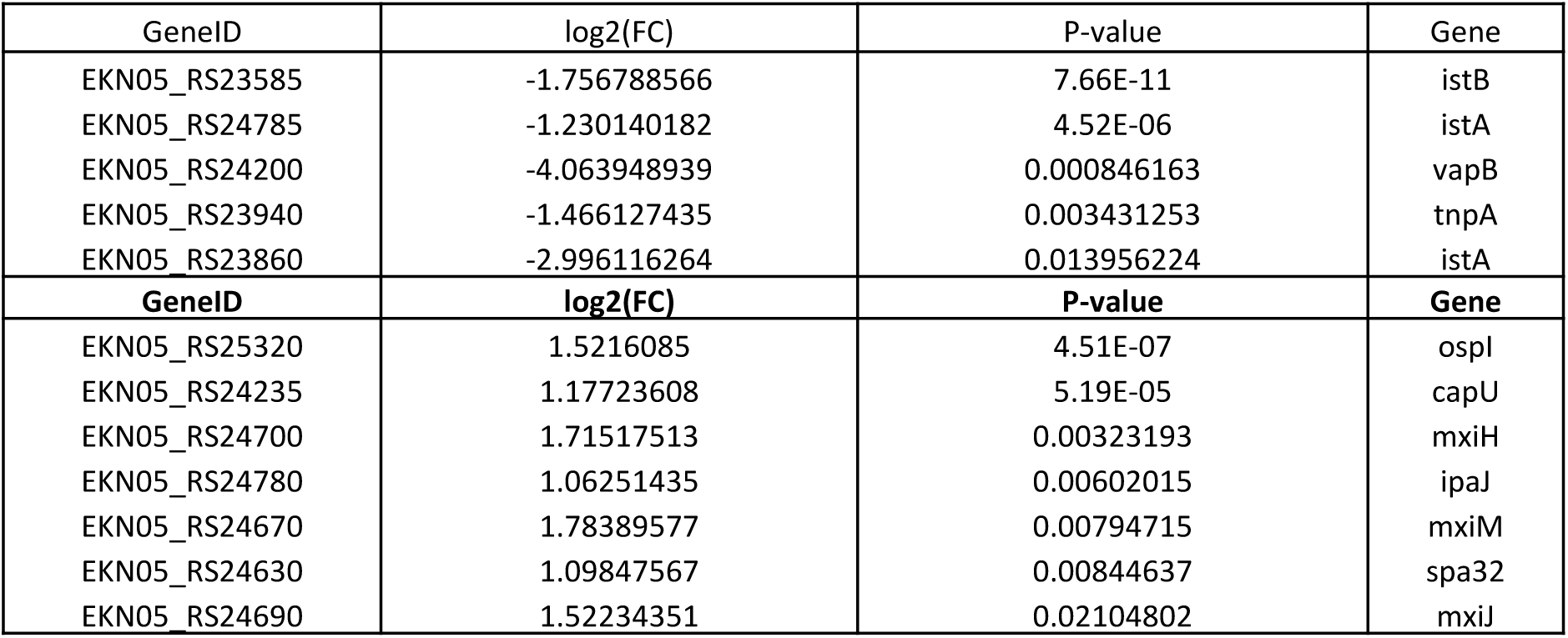
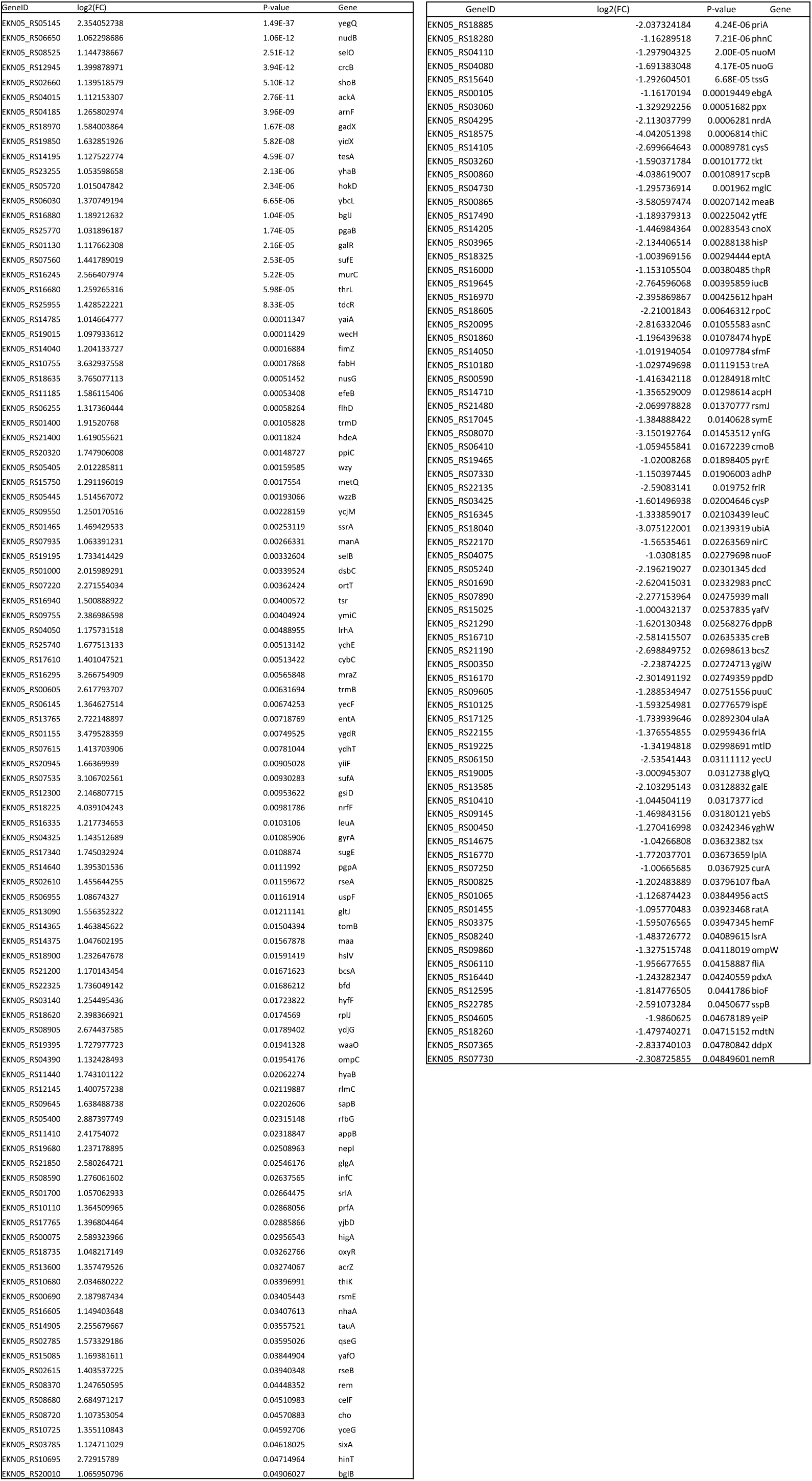

**Table S3.**

| s | GS<br> follow link to MSigDB | GS DETAILS | SIZE | ES | NES | NOM p-val | FDR q-val | FWER p-val | RANK AT MAX | LEADING EDGE |
| --- | --- | --- | --- | --- | --- | --- | --- | --- | --- | --- |
| HALLMARK_INTERFERON_ALPHA_RESPONSE | HALLMARK_INTERFERON_ALPHA_RESPONSE | Details ... | 97 | 0.7108243 | 2.2621415 | 0.0 | 0.0 | 0.0 | 3627 | tags=48%, list=12%, signal=55% |
| HALLMARK_INTERFERON_GAMMA_RESPONSE | HALLMARK_INTERFERON_GAMMA_RESPONSE | Details ... | 200 | 0.593273 | 2.0486922 | 0.0 | 0.0 | 0.0 | 4615 | tags=41%, list=15%, signal=48% |
| HALLMARK_INFLAMMATORY_RESPONSE | HALLMARK_INFLAMMATORY_RESPONSE | Details ... | 200 | 0.42803988 | 1.4914694 | 0.0 | 0.05034907 | 0.118 | 5374 | tags=36%, list=18%, signal=43% |
| HALLMARK_P53_PATHWAY | HALLMARK_P53_PATHWAY | Details ... | 200 | 0.41123477 | 1.4398199 | 0.0 | 0.070149854 | 0.204 | 7187 | tags=46%, list=24%, signal=60% |
| HALLMARK_CHOLESTEROL_HOMEOSTASIS | HALLMARK_CHOLESTEROL_HOMEOSTASIS | Details ... | 74 | 0.47468454 | 1.4319116 | 0.012345679 | 0.06316165 | 0.227 | 6999 | tags=47%, list=23%, signal=61% |
| HALLMARK_ALLOGRAFT_REJECTION | HALLMARK_ALLOGRAFT_REJECTION | Details ... | 200 | 0.40094516 | 1.4020487 | 0.0033167496 | 0.073521934 | 0.306 | 5597 | tags=28%, list=19%, signal=34% |
| HALLMARK_IL6_JAK_STAT3_SIGNALING | HALLMARK_IL6_JAK_STAT3_SIGNALING | Details ... | 87 | 0.435721 | 1.3675903 | 0.02640845 | 0.09310555 | 0.422 | 4660 | tags=31%, list=15%, signal=37% |
| HALLMARK_HYPOXIA | HALLMARK_HYPOXIA | Details ... | 200 | 0.38327393 | 1.3310933 | 0.009708738 | 0.12024634 | 0.567 | 7245 | tags=43%, list=24%, signal=56% |
| HALLMARK_KRAS_SIGNALING_UP | HALLMARK_KRAS_SIGNALING_UP | Details ... | 200 | 0.37829182 | 1.3203443 | 0.0121951215 | 0.12094211 | 0.619 | 6585 | tags=38%, list=22%, signal=48% |
| HALLMARK_EPITHELIAL_MESENCHYMAL_TRANSITION | HALLMARK_EPITHELIAL_MESENCHYMAL_TRANSITION | Details ... | 200 | 0.36944866 | 1.2812766 | 0.033557046 | 0.16590218 | 0.769 | 3572 | tags=23%, list=12%, signal=26% |
| HALLMARK_APICAL_JUNCTION | HALLMARK_APICAL_JUNCTION | Details ... | 200 | 0.36928305 | 1.2758917 | 0.028145695 | 0.15843831 | 0.789 | 7026 | tags=40%, list=23%, signal=51% |
| HALLMARK_MYOGENESIS | HALLMARK_MYOGENESIS | Details ... | 200 | 0.3657497 | 1.2716538 | 0.029173419 | 0.15033501 | 0.804 | 6957 | tags=36%, list=23%, signal=47% |
| HALLMARK_COAGULATION | HALLMARK_COAGULATION | Details ... | 138 | 0.37270144 | 1.245964 | 0.07781457 | 0.17971285 | 0.877 | 6400 | tags=30%, list=21%, signal=38% |
| HALLMARK_ESTROGEN_RESPONSE_LATE | HALLMARK_ESTROGEN_RESPONSE_LATE | Details ... | 200 | 0.35684922 | 1.2419971 | 0.040472176 | 0.1734811 | 0.886 | 3669 | tags=25%, list=12%, signal=28% |
| HALLMARK_XENOBIOTIC_METABOLISM | HALLMARK_XENOBIOTIC_METABOLISM | Details ... | 200 | 0.3534178 | 1.2355049 | 0.0544 | 0.17134531 | 0.898 | 6202 | tags=33%, list=21%, signal=41% |
| HALLMARK_HEDGEHOG_SIGNALING | HALLMARK_HEDGEHOG_SIGNALING | Details ... | 36 | 0.44858375 | 1.2176301 | 0.16098484 | 0.1889897 | 0.935 | 4991 | tags=36%, list=17%, signal=43% |
| HALLMARK_KRAS_SIGNALING_DN | HALLMARK_KRAS_SIGNALING_DN | Details ... | 200 | 0.34908617 | 1.211161 | 0.06456953 | 0.18889388 | 0.944 | 5875 | tags=29%, list=20%, signal=35% |
| HALLMARK_IL2_STAT5_SIGNALING | HALLMARK_IL2_STAT5_SIGNALING | Details ... | 199 | 0.34493807 | 1.1918249 | 0.09059829 | 0.21274173 | 0.97 | 3729 | tags=21%, list=12%, signal=24% |
| HALLMARK_COMPLEMENT | HALLMARK_COMPLEMENT | Details ... | 200 | 0.3271525 | 1.1394963 | 0.13483146 | 0.31150877 | 0.995 | 6610 | tags=35%, list=22%, signal=44% |
| HALLMARK_ANGIOGENESIS | HALLMARK_ANGIOGENESIS | Details ... | 36 | 0.41598716 | 1.116682 | 0.26725665 | 0.35646445 | 0.998 | 3643 | tags=25%, list=12%, signal=28% |
| HALLMARK_APOPTOSIS | HALLMARK_APOPTOSIS | Details ... | 161 | 0.32843515 | 1.0973455 | 0.23076923 | 0.39179388 | 1.0 | 7224 | tags=38%, list=24%, signal=50% |
| HALLMARK_ADIPOGENESIS | HALLMARK_ADIPOGENESIS | Details ... | 200 | 0.3095652 | 1.0756397 | 0.25566342 | 0.43653414 | 1.0 | 6969 | tags=37%, list=23%, signal=48% |
| HALLMARK_OXIDATIVE_PHOSPHORYLATION | HALLMARK_OXIDATIVE_PHOSPHORYLATION | Details ... | 200 | 0.30664378 | 1.0641147 | 0.2677966 | 0.45221516 | 1.0 | 7071 | tags=35%, list=23%, signal=45% |
| HALLMARK_BILE_ACID_METABOLISM | HALLMARK_BILE_ACID_METABOLISM | Details ... | 112 | 0.3074014 | 0.99455786 | 0.47386172 | 0.6680501 | 1.0 | 6903 | tags=33%, list=23%, signal=43% |
| HALLMARK_GLYCOLYSIS | HALLMARK_GLYCOLYSIS | Details ... | 200 | 0.2803198 | 0.978534 | 0.5247209 | 0.7014511 | 1.0 | 6322 | tags=29%, list=21%, signal=37% |
| HALLMARK_REACTIVE_OXYGEN_SPECIES_PATHWAY | HALLMARK_REACTIVE_OXYGEN_SPECIES_PATHWAY | Details ... | 49 | 0.33910406 | 0.9722521 | 0.5138632 | 0.69652385 | 1.0 | 5892 | tags=33%, list=20%, signal=41% |
| HALLMARK_PEROXISOME | HALLMARK_PEROXISOME | Details ... | 104 | 0.2997349 | 0.95374775 | 0.5533808 | 0.73891777 | 1.0 | 6934 | tags=35%, list=23%, signal=45% |
| HALLMARK_HEME_METABOLISM | HALLMARK_HEME_METABOLISM | Details ... | 200 | 0.2704876 | 0.9417018 | 0.622549 | 0.7543854 | 1.0 | 4614 | tags=23%, list=15%, signal=27% |
| HALLMARK_ESTROGEN_RESPONSE_EARLY | HALLMARK_ESTROGEN_RESPONSE_EARLY | Details ... | 200 | 0.27311674 | 0.9412839 | 0.6365132 | 0.72974205 | 1.0 | 4587 | tags=25%, list=15%, signal=29% |
| HALLMARK_UNFOLDED_PROTEIN_RESPONSE | HALLMARK_UNFOLDED_PROTEIN_RESPONSE | Details ... | 113 | 0.29046243 | 0.92753977 | 0.6167247 | 0.75246346 | 1.0 | 4125 | tags=22%, list=14%, signal=26% |
| HALLMARK_TNFA_SIGNALING_VIA_NFKB | HALLMARK_TNFA_SIGNALING_VIA_NFKB | Details ... | 200 | 0.2679294 | 0.9219561 | 0.68604654 | 0.74633217 | 1.0 | 3647 | tags=25%, list=12%, signal=28% |
| HALLMARK_ANDROGEN_RESPONSE | HALLMARK_ANDROGEN_RESPONSE | Details ... | 101 | 0.2767761 | 0.89216363 | 0.73905724 | 0.8181602 | 1.0 | 6364 | tags=30%, list=21%, signal=38% |
| HALLMARK_PI3K_AKT_MTOR_SIGNALING | HALLMARK_PI3K_AKT_MTOR_SIGNALING | Details ... | 105 | 0.24169427 | 0.7779215 | 0.95555556 | 1.0 | 1.0 | 7062 | tags=31%, list=23%, signal=41% |
| HALLMARK_UV_RESPONSE_UP | HALLMARK_UV_RESPONSE_UP | Details ... | 158 | 0.22651502 | 0.7590925 | 0.9933665 | 1.0 | 1.0 | 7175 | tags=35%, list=24%, signal=45% |
| HALLMARK_APICAL_SURFACE | HALLMARK_APICAL_SURFACE | Details ... | 44 | 0.22457846 | 0.6259953 | 1.0 | 0.99949026 | 1.0 | 7286 | tags=27%, list=24%, signal=36% |

**Table S4.**
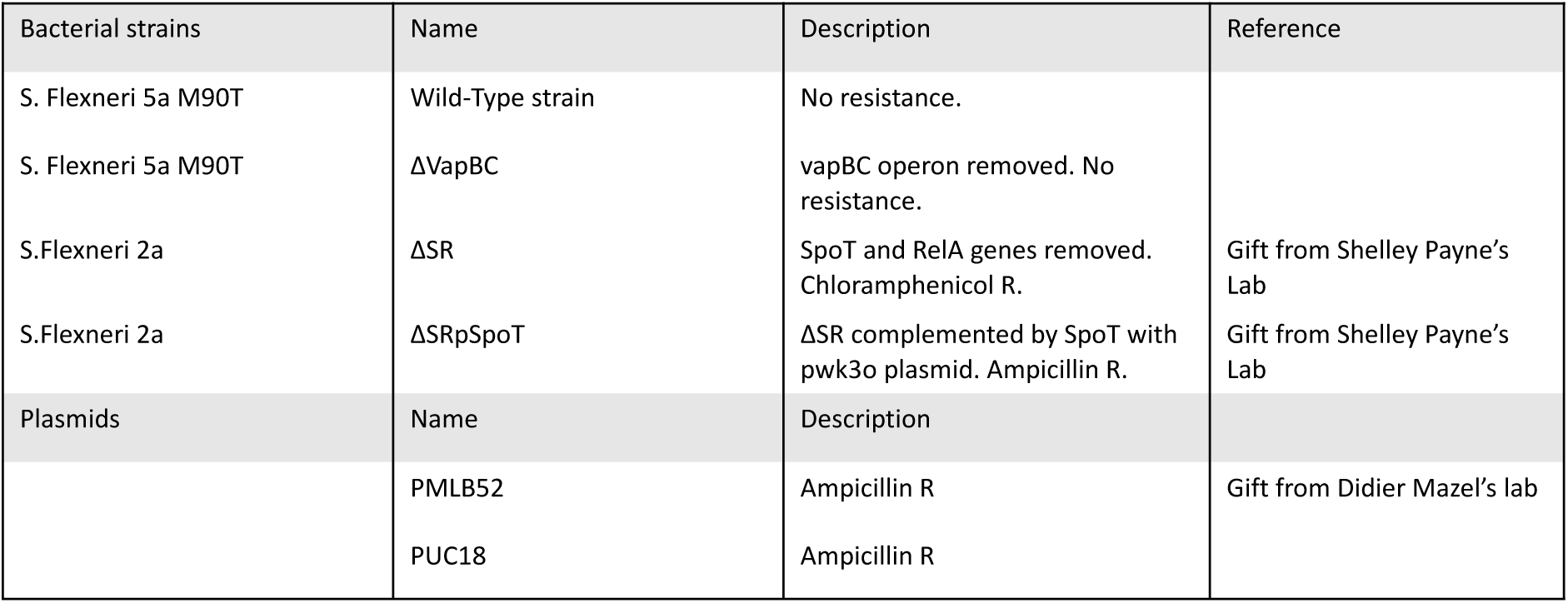
List of the bacterial strains and plasmids used in this study.

**Table S5.**
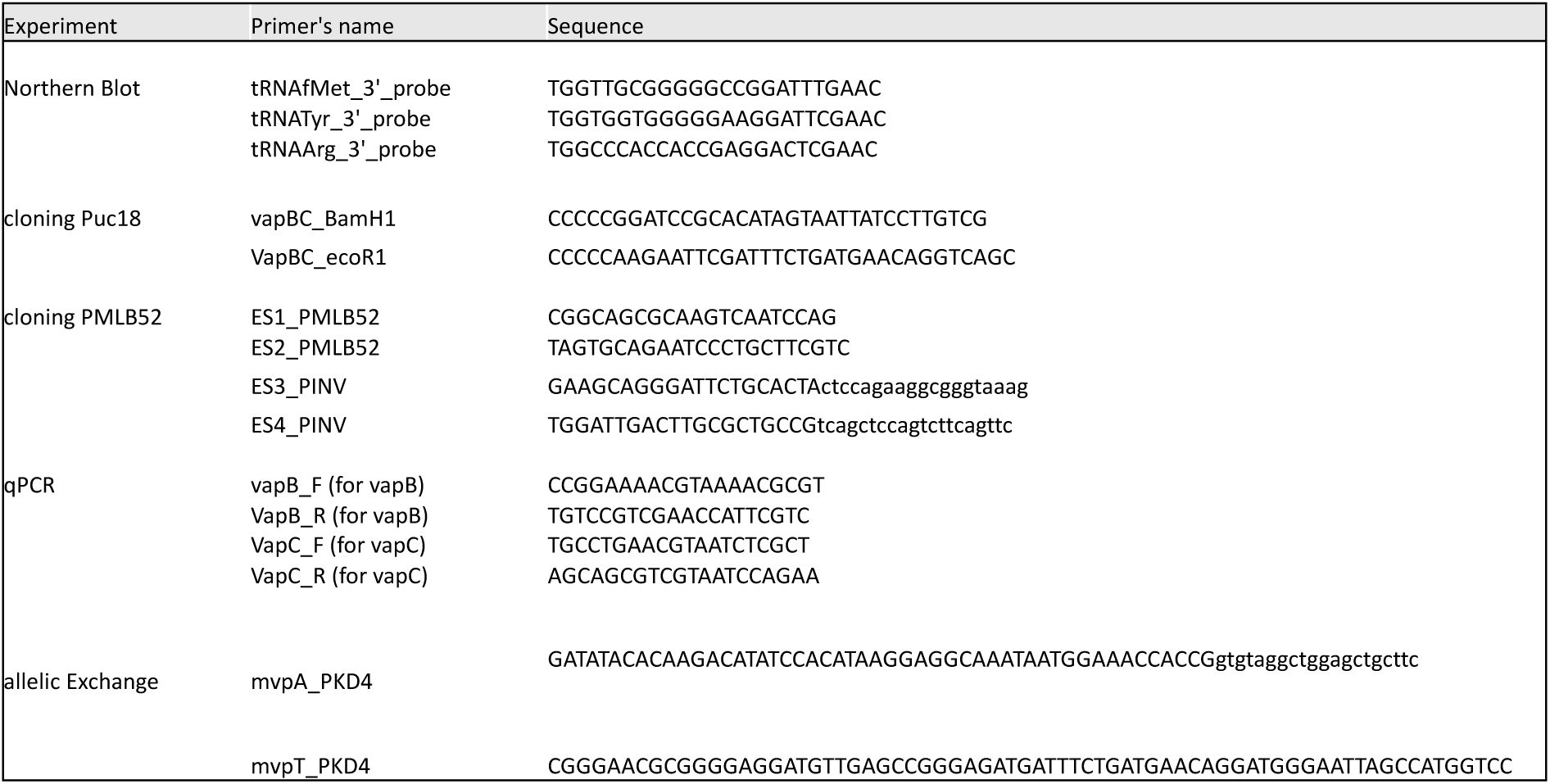
List of primers used in this study.

## Material and methods

### Bacterial Strains and plasmid

The strains and plasmids and primers used in this study are listed in the **Supplementary Table S4 and S5.** Bacteria were grown in TSB broth or TSB agar plates. All mutants were constructed using the λ red linear recombination method as described using the primer describes in **Supplementary S5** (Datsenko & Wanner, 2000). For low copy plasmid complemented strains, *vapBC* was amplified by PCR using Phusion DNA polymerase (Thermo Fisher Scientific, F-530L). Following amplification, 1 µL of DpnI enzyme (Thermo Fisher Scientific, FD1704) was added directly to the PCR product to eliminate the methylated parental template DNA, and the reaction was incubated at 37°C for 1 hour. The resulting DNA fragment was purified using the Macherey-Nagel NucleoSpin Gel and PCR Clean-Up Kit (REF 740609). The low-copy plasmid PMLB52 (a generous gift from Didier Mazel, Pasteur Institute) was used, and the construct was assembled using the Gibson Assembly method with HiFi DNA Assembly Mix (New England Biolabs, E2621). For the high-copy plasmid pUC18 was used. Both the vector and PCR product were digested with BamHI (NEB #R3101) and EcoRI (NEB #R3101) at 37 °C for 1 hour, and the resulting fragments were purified using the same Macherey-Nagel kit (REF 740609.50).

### Production of Anti-VapB and Anti-VapC Rabbit Polyclonal Antibodies

Recombinant VapB and VapC proteins were used to immunize two rabbits (Cusabio Technology LLC). Total serum was collected from both animals one month after immunization and tested by western blot for specific recognition of the purified proteins as well as the native proteins in *Shigella* lysates. The resulting antisera showed specific detection at a 1:2000 dilution in western blotting and were subsequently used for further experiments.

### Cell Culture

The enterocytic cell line HCT116 was a gift from B. Vogelstein (Johns Hopkins University, Baltimore, MD, USA). These cells were all grown in Dulbecco’s modified Eagle’s medium (DMEM) supplemented with 10% FBS.

### Bacterial infection in epithelial cells

Overnight bacterial cultures were incubated at 30 °C with shaking at 160 rpm, then diluted 1:35 and further grown at 37 °C with agitation (160 rpm) until reaching an optical density (OD) of 0.7. The bacterial cells were pelleted and resuspended in serum-free DMEM. This suspension was added to host cells that had been serum-starved for 30 minutes and had reached ∼70% confluency. Infections were carried out at a multiplicity of infection (MOI) of 100 for 5 hours of infection and 0.5 for 16 hours of infection. The culture plates were centrifuged at room temperature for 10 minutes at 2000 rpm to synchronize infection. Subsequently, plates were incubated for 30 minutes under humidified conditions at 37 °C and 5% CO₂. After incubation, the cells were washed three times with PBS, and fresh serum-free DMEM containing 50 µg/mL gentamicin was added. This point was designated as time zero of infection.

### T3SS secretion assay

The Shigella strains were cultured until the late exponential phase in 100 mL of TSB broth. The bacterial density was normalized by measuring the optical density (OD). Bacteria were harvested by centrifugation for 10 minutes at 9500 rpm, resuspended in 4 mL of PBS, and divided into two 2 mL samples. Each sample was either supplemented with 100 μg/mL of Congo Red or left untreated and then incubated for 30 minutes at 37°C to induce the type III secretion system (TTSS). Subsequently, bacteria were collected again by centrifugation for 10 minutes at 10,000 rpm. A fraction of the pellet (1/150th) was suspended in Laemmli buffer, while the supernatant was precipitated with trichloroacetic acid (TCA) and then suspended in Laemmli buffer.

### Congo Red (CR) binding assay

Bacterial cultures at the indicated times were streaked onto Tryptic Soy Agar (TSA) plates supplemented with 0.01% (w/v) Congo Red dye (Sigma-Aldrich). The plates were incubated at 37°C for 24 hours. Post-incubation, colony coloration was evaluated. Virulent strains retaining the pINV plasmid formed red or pink colonies due to Congo Red binding, whereas avirulent or plasmid-cured strains produced white or colorless colonies. This phenotypic distinction served as an indicator of plasmid pINV presence.

### FC Analyses of TSAR-Expressing Bacteria Recovered from Infected Cells

HCT116 cells were infected at MOI50 during 5 hours with bacteria harboring the pTSAR 1.3 plasmid (Campbell-Valois et al., 2014). Intracellular bacteria were recovered as described(Aussel et al., 2011), and subjected to FC analyses for analyzing T3SS activation.

### Iron starvation

*Shigella flexneri* strains were grown overnight in BTCS broth at 37°C with shaking. The following day, cultures were diluted into fresh BTCS medium supplemented with 0.25 mM 2,2ʹ-dipyridyl (an iron chelator dissolved in ethanol) and incubated at the indicated times at 37°C with shaking to induce iron limitation. Control cultures were treated with an equivalent volume of ethanol to account for any solvent effects. Following incubation, bacterial cells were collected by centrifugation and pellets were processed for downstream analyses, including Western blotting to assess protein expression and Northern blotting to evaluate RNA profiles under iron-depleted conditions.

### Northern Blot Analysis

The bacterial pellet was resuspended in 1 ml of Trizol reagent and homogenized by pipetting up and down to facilitate RNA extraction. RNA extraction followed the guidelines for RNA isolation using TRIzol™ Reagent (Invitrogen, Cat. No. 15596026).

Total RNA (10 µg) was extracted using the Nucleospin miRNA kit (Macherey-Nagel) to separate small RNAs (<200 nt). Five micrograms of small RNA per sample were resolved on a 12% acrylamide/8 M urea/MOPS denaturing gel and run at 300 V. After electrophoresis, RNA loading was assessed by BET staining under UV illumination. RNAs were transferred to a Hybond N⁺ membrane (Amersham) in 1× MOPS buffer at 20 V for 1 h at 4 °C and crosslinked using EDC (1-ethyl-3-(3-dimethylaminopropyl) carbodiimide hydrochloride).

DNA probes **(sequence in Table S5)** were 5ʹ end-labeled with [γ-³²P] ATP using T4 PNK (New England Biolabs), purified on G-25 spin columns, and denatured at 95 °C for 5 min. Hybridization was carried out overnight at 40 °C in UltraHyb-Oligo buffer (Invitrogen). Membranes were washed sequentially with 2× SSC/0.5% SDS and 1× SSC/0.5% SDS

### Dual RNA Sequencing (Dual RNA-seq)

HCT116 cells were infected with WT or Δ*vapBC Shigella* strains at MOI 100. At 5 hours or 16 hours post-infection, total RNA from infected monolayers was extracted using TRIzol and treated with DNase. rRNA was depleted using Ribo-Zero (Illumina) and libraries were prepared using the TruSeq Stranded Total RNA Library Prep Kit (Illumina). Sequencing was performed on an Illumina platform. Differential gene expression was analyzed using DESeq2. Genes with an adjusted p-value < 0.05 and log₂ fold change ≥ |1| were considered significant.

### qRT–PCR

The *Shigella* culture grown under anaerobic conditions for 2.5 hours was harvested by centrifugation at 6000 rpm for 10 minutes at 37°C for subsequent qRT–PCR analysis. The resulting pellet was resuspended in 1 ml of Trizol reagent and homogenized by pipetting up and down to facilitate RNA extraction. RNA extraction followed the guidelines for RNA isolation using TRIzol™ Reagent (Invitrogen, Cat. No. 15596026). To eliminate potential genomic DNA contamination in the RNA sample, Turbo DNase I (Invitrogen, AM2238) digestion was performed. Subsequently, the RNA underwent a purification step using acidic phenol/chloroform extraction procedures. The purified RNA was then reverse transcribed to synthesize cDNA using the RevertAid H Minus First Strand cDNA Synthesis Kit (Thermo Scientific, K1632) following the manufacturer’s recommendations. For qPCR, the Brilliant III Ultra-Fast SYBR Green QPCR Master Mix (Agilent, 600882) was employed.

### Western blot

Bacterial pellets were denatured by adding Laemmli sample buffer supplemented with β-mercaptoethanol, followed by incubation at 95 °C for 5 minutes with shaking. Protein separation was performed using 15% Tris/Glycine SDS-PAGE gels (1.5 mm thick). Proteins were then transferred to nitrocellulose membranes using a wet transfer system with Tris/Glycine buffer containing 15% methanol, at 90 V for 1 hour and 30 minutes at 4 °C. Membranes were blocked for 30 minutes in TBST containing 5% non-fat dry milk. After a brief rinse in TBST, membranes were incubated overnight at 4 °C with gentle agitation in primary antibody diluted in TBST. For detection of VapB and VapC, custom-made or Cusabio anti-VapB/anti-VapC antibodies were used at a dilution of 1:200 in 5% milk/TBST following prior antibody depletion. The anti-RecA antibody (Abcam, ab63797) was diluted 1:1000 in 5% milk/TBST. Secondary detection was performed using an anti-rabbit HRP-conjugated antibody (MP Biomedicals, 08674371) diluted 1:5000 in 5% milk/TBST, incubated for 1 hour at room temperature.

### Plaque assay

The protocol used is from Reisacher et al (2025) (Reisacher et al., 2025). Bacterial infection was carried out on 100% confluent cells at a multiplicity of infection of 0.5. Cells with bacterial suspension were incubated for 2h in a humid atmosphere, 37°C temperature and 5% CO2. Cells were then washed three times with PBS. Then, DMEM with 10% fetal calf serum, and supplemented with 50 µg/ml Gentamicin, was added to the cells. Plates were incubated for 24h in a humid atmosphere, 37°C temperature and 5% CO2. Cells were then fixed and stained with DAPI for imaging.

### RNA extraction

Total RNA was extracted using TRIzol reagent (Life Technologies, 15596026), following the manufacturer’s protocol, at a ratio of 1 mL TRIzol per 6 million cells or fewer. Briefly, cells were lysed directly in the culture dish by adding TRIzol and incubating for 5 minutes at room temperature. Phase separation was achieved by adding 200 µL of chloroform (Sigma-Aldrich, 372978), followed by centrifugation at 12,000 × g for 15 minutes at 4 °C. The aqueous phase was carefully collected, and an equal volume of isopropanol (Sigma-Aldrich, I9516) was added to precipitate RNA. After homogenization, samples were incubated at −20 °C overnight. Precipitated RNA was pelleted by centrifugation at 20,000 × g for over 30 minutes at 4 °C, then washed twice with ice-cold 70% nuclease-free ethanol (Sigma-Aldrich, 51976). The RNA pellet was briefly air-dried at room temperature and resuspended in nuclease-free water (Life Technologies, 10977035). RNA samples were stored at −80 °C until further use.

### Microscopy image acquisition and quantification

Microscopy Images were acquired on a structured illumination fluorescent microscope Apotome 2 (Zeiss). Fluorescence quantification was performed using ImageJ software (Rasband, 1997-2018).

### Timelapse imaging

Bacterial cultures were grown overnight at 30°C, and inoculated at a 1:100 dilution in TCSB supplemented with the appropriate antibiotics, and grown to an optical density between 0.4 and 0.6 at 600 nm (OD600). For time lapse imaging experiments, prior to infection, bacteria were washed twice with EM buffer (120 mM NaCl, 7 mM KCl, 1.8 mM CaCl2, 0.8 mM MgCl2, 5 mM glucose, 25 mM HEPES, pH 7.3) and finally diluted in EM buffer to reach a MOI of 20. Before adding bacteria to the target cells, they were coated with poly-L-lysine (10 μg/mL) for 10 min at 37°C and washed twice in EM buffer prior to dilution to reach a MOI of 100. In 8 well-Ibidi chambers (clinisciences), 100 μL per well of the diluted bacterial suspension was added to the cells.

Infected HeLa cells stably expressing galectin-3-eGFP(Fredlund et al., 2018) in EM buffer were imaged at 37°C using a spinning disc (CSU-W1) confocal microscope (Nikon) using a 40X/0.75NA air objective. Cells were imaged every 2 minutes for 120 min, on a range of 8 μm, with a section spacing of 0.3 μm in the Z dimension and using a piezo control and the UltimateFocus module to keep the plane in focus. FITC excitation and emission filters were used to measure the eGFP signals. Image analysis was systematically performed using Fiji (http://fiji.sc). For BCV unpeeling measurements, maximum Z-projections were used. The galectin-3 signal was used to measure the onset of vacuolar rupture at the timepoint of prominent recruitment, and to analyze the onset of unpeeling when the signal started to change morphology becoming separated from the entering bacteria. Changing the fluorescent parameters also enabled the start of bacterial entry with the galectin-3 signal due to the ruffle formation at the entry site. Quantification of unpeeling was done as described before(Mellouk et al., 2024b).

### Statistical analysis

Statistical analysis was performed using GraphPad Prism 8.0.1

## Acknowledgements

We thank the imaging and cytometry platforms at the SFR Necker and Nathalie Servel of the radioactivity core facility of the INEM for their technical help. This work was supported by the INSERM dotation. E. Saifi was supported by the BioSPC doctoral school and by the funding program of the G.E.N.E. Graduate School of Université Paris Cité.

